# Age-dependent Effects of Titrating Anodal Transcranial Direct Current Stimulation (tDCS) Intensity on Motor Sequence Learning

**DOI:** 10.64898/2026.06.09.731079

**Authors:** Amba M. Frese, Ruxandra Ungureanu, Ensiyeh Ghasemian-Shirvan, Lorena Melo, Yiwu Xiong, Marie C. Beaupain, Min-Fang Kuo, Raf L. J. Meesen, Michael A. Nitsche

## Abstract

Optimising the currently heterogenous efficacy of transcranial direct current stimulation (tDCS) interventions for motor learning requires identifying stimulation parameters facilitating performance while accounting for age-related differences in baseline performance and mechanisms underlying neuroplasticity.

We systematically explored anodal tDCS intensity effects on implicit motor sequence learning (IMSL) in young and older adults. The study utilised a randomised, double-blind, counterbalanced crossover design. Ninety-six healthy participants (48 young adults, 48 older adults) completed a serial reaction time task (SRTT) with online sham or anodal tDCS over M1 at intensities of 1, 2, and 3 mA. The next day, memory consolidation was assessed in a recall test.

Both age groups demonstrated IMSL and consolidation across conditions. While 1 mA tDCS improved IMSL in young adults by reducing reaction times, higher intensities had no significant benefit compared to sham. In older adults, anodal tDCS did not affect general task performance compared to sham, but 1mA tDCS acutely impaired selective sequence learning.

The results demonstrate age-dependent and non-linear dose-dependent effects of anodal tDCS on IMSL. This underscores the necessity for age-adapted protocols for experimental and clinical tDCS applications. Future research should explore neurophysiological reasons for reduced tDCS efficacy in older adults found in the present study.

## 1. Introduction

Aging is associated with reduced neuroplasticity, impairing cognitive and motor functions in later life [1–3]. This is particularly relevant for implicit motor sequence learning (IMSL), essential for acquiring everyday sequential skills and maintaining independence in older age [4]. Transcranial direct current stimulation (tDCS) non-invasively induces plasticity and enhances IMSL, but age-related differences in tDCS parameter effects on IMSL remain underexplored.

During tDCS, targeted neural populations are de- or hyperpolarised at a subthreshold level [5]. Key parameters affecting tDCS efficacy include current density and intensity, electrode size and montage, and stimulation polarity [5]. Anodal tDCS after-effects resemble long-term potentiation (LTP), increasing glutamatergic NMDA receptor (NMDAR) activity and reducing GABAergic inhibition [6–8]. The parameter space of anodal tDCS has largely been explored for primary motor cortex (M1) excitability. Protocol-dependent facilitation patterns with stimulation intensities between 1-3 mA and 15-30 min durations have shown non-linear effects in young adults, with minor differences between conditions. In contrast, lower intensities (1 mA) and relatively short stimulation durations showed decreased facilitation in older adults, aligned with higher age-related LTP-like plasticity induction thresholds [9–11]. To translate tDCS intensity parameters to cognitive outcomes, such as IMSL, studies titrating tDCS intensities and exploring specific effects on cognition are required.

In IMSL, novel movement sequences are acquired without conscious awareness or intent [12–14]. Aging results in slower reaction times in IMSL tasks, alongside impaired consolidation and retention [15–17]. Anodal tDCS has shown facilitation of motor functions in healthy young and older adults, as well as patients with motor deficits or age-related neurological conditions [4,18–24]. Although anodal tDCS over M1 enhances IMSL and consolidation, the optimal stimulation intensity remains unclear [4,25,26]. Cuypers and colleagues (2013) found that 1.5 mA anodal M1 tDCS improved motor sequence learning relative to sham in young adults, whereas 1mA did not [20]. However, other studies found that 10-20 min anodal tDCS over M1 with 1-2 mA intensity facilitated motor learning and consolidation in young adults [4,27–31]. Results are somewhat heterogeneous, partly due to homeostatic tDCS and learning interactions, where online tDCS enhances motor sequence learning while pre-task stimulation impairs it [30,32,33]. Findings in older adults are similarly mixed, with some reporting no effects of anodal tDCS intensities of 1-3 mA on IMSL, and others finding improvements in motor performance with 1 mA [21–24,34–37]. Together, this highlights the need for systematic investigations of how tDCS intensity influences IMSL across age groups.

Age-related neurophysiological changes challenge tDCS efficacy in older adults. In aging, structural volumetric changes in the head, skin, and CSF reduce received tDCS-induced electrical field strength and alter current distribution, reducing tDCS effects and increasing inter-individual variability [38,39]. Additionally, the acquisition of motor memories involves the strengthening of motor learning related synapses primarily through glutamatergic LTP, gated by GABAergic effects [40]. Correspondingly, across age groups, motor learning magnitude positively correlates with learning-related decreases in GABA, disinhibiting glutamatergic activation [7,35,41–43]. This excitatory/ inhibitory (E/I) balance of glutamate and GABA is altered with normal aging, due to decreased GABA concentrations and NMDAR availability and function, which limit E/I changes responding to task demands [44–48]. These age-related E/I alterations are associated with impairments in motor performance [48–50]. Because anodal tDCS reduces GABA, age-related GABA decreases may limit its effects. An MRS study found that both motor learning alone and anodal tDCS post-learning did not reduce GABA levels in the sensorimotor cortex of elderly participants, suggesting a floor effect [35]. However, only 1mA tDCS was applied in that study, which may be insufficient given the age-related attenuation of induced electrical field strength.

Currently, no study has systematically titrated tDCS intensities while assessing IMSL and comparing effects between age groups. Here, we titrated anodal tDCS intensities (1mA, 2mA, 3mA, Sham) over M1 of both young and older adults during an IMSL task (SRTT) with an additional next-day assessment of consolidation. We hypothesised that older adults would still show IMSL, but reduced consolidation compared to young adults. Additionally, we hypothesised that anodal tDCS would result in a non-linear, dosage-dependent facilitation of IMSL in young adults, with young adults benefitting more from lower tDCS intensities than older adults. Older adults were hypothesised to benefit less from anodal tDCS than young adults, due to weaker tDCS-induced electrical fields.

## 2. Results

One elderly participant was excluded due to <50% SRTT accuracy. Outliers removed per block averaged 1.3 ± 0.9% for young and 1.5 ± 0.9% for elderly participants (mean ± SD). After all experimental sessions, 29 young and 40 elderly participants reported noticing a repeating sequence.

### 2.1. Learning: Effects of Age and tDCS Intensity on SRTT Performance during tDCS

For absolute RTs, the ANOVA showed significant main and interaction effects of Block, Age Group, Block * Age Group, and tDCS Condition * Block * Age Group. Other main and interaction effects were not significant (Table 1). Post-hoc comparisons revealed that both young and elderly samples had significantly faster RTs in sequence blocks (blocks 2-5, 7-8) compared to the respective block 1 across tDCS conditions. In young adults, RTs in block 6 with 2mA and 3mA tDCS were significantly slower than block 1. Elderly participants had significantly slower RTs than young adults across blocks and tDCS conditions. The young sample’s RTs in the 1mA tDCS condition were significantly faster than sham at blocks 2-5 and 8. The elderly sample’s RTs did not differ significantly between active and sham tDCS (Figure 2).

**Table 1:** Results of the Mixed Model ANOVAs of Absolute Reaction Time in the Serial Reaction Time Task (SRTT). Mauchly’s test of sphericity was conducted for each ANOVA, and Greenhouse–Geisser corrections were applied where sphericity assumptions were not met. Asterisks indicate significant effects (p<.05). d.f. = degrees of freedom; n2p = partial eta squared.

| *Variables* | *d.f.* | *d.f.error* | *F-value* | *n2p* | *p-value* |
| --- | --- | --- | --- | --- | --- |
| **Learning (during tDCS)** |  |  |  |  |  |
| tDCS Condition | 3 | 279 | .417 | .004 | .741 |
| Block | 1.625 | 151.122 | 63.991 | .408 | **<.001\*** |
| Age Group | 1 | 93 | 49.629 | .348 | **<.001\*** |
| tDCS Condition X Block | 10.334 | 961.062 | .670 | .007 | .758 |
| tDCS Condition X Age Group | 3 | 279 | 1.569 | .017 | .197 |
| Block X Age Group | 1.625 | 151.122 | 16.090 | .147 | **<.001\*** |
| tDCS Condition X Block X Age Group | 10.334 | 961.062 | 1.876 | .020 | **.043\*** |
| **Recall (next day)** |  |  |  |  |  |
| tDCS Condition | 3 | 279 | .807 | .009 | .491 |
| Block | 1.104 | 102.698 | 96.717 | .510 | **<.001\*** |
| Age Group | 1 | 93 | 51.649 | .357 | **<.001\*** |
| tDCS Condition X Block | 4.242 | 394.547 | 2.450 | .026 | **.042\*** |
| tDCS Condition X Age Group | 3 | 279 | 1.228 | .013 | .300 |
| Block X Age Group | 1.104 | 102.698 | 19.074 | .170 | **<.001\*** |
| tDCS Condition X Block X Age Group | 4.242 | 394.547 | 2.062 | .022 | .081 |

For baseline-standardised RTs, the ANOVA showed significant main and interaction effects of Block, Age Group, Block * Age Group, and tDCS Condition * Block * Age Group. No other main or interaction effects were significant (Table S1.1). Post-hoc comparisons revealed significantly faster baseline-standardised RTs in all sequence blocks compared to baseline across tDCS conditions for both groups. In young participants, RTs in block 6 were significantly slower than baseline with 1mA tDCS. In elderly participants, RTs in block 6 were significantly slower than baseline with sham and 2mA tDCS. Across tDCS conditions, the elderly sample had significantly slower baseline-standardised RTs in sequence blocks than the young sample. Within the young sample, baseline-standardised RTs with 1mA tDCS were significantly faster in blocks 2-5 and 8 compared to sham (Figure S2.1).

For selective sequence learning (blocks 5 & 6), the ANOVA showed significant main and interaction effects of Block, Age Group, Block * Age Group, and tDCS Condition * Block * Age Group. Remaining main and interaction effects were not significant (Table 2). Post hoc tests for the difference in block 5 and 6 baseline-standardised RTs (Equation 1) showed that older adults had significantly lower selective sequence learning than young adults across tDCS conditions. Compared to sham, 1mA tDCS significantly increased selective sequence learning in young adults but significantly reduced it in older adults (Figure 2).

**Table 2:** Results of the Mixed Model ANOVA of the Main Selective Sequence Learning Blocks 5 and 6 in the Serial Reaction Time Task (SRTT). Mauchly’s test of sphericity was conducted, confirming that sphericity assumptions were met. Asterisks indicate significant effects (p<.05). d.f. = degrees of freedom; n2p = partial eta squared.

| *Variables* | *d.f.* | *d.f.<sub>error</sub>* | *F-value* | *n2p* | *p-value* |
| --- | --- | --- | --- | --- | --- |
| tDCS Condition | 3 | 279 | .083 | .001 | 0.969 |
| Block | 1 | 93 | 100.911 | .520 | **<.001\*** |
| Age Group | 1 | 93 | 43.045 | .316 | **<.001\*** |
| tDCS Condition X Block | 3 | 279 | .266 | .003 | 0.850 |
| tDCS Condition X Age Group | 3 | 279 | .853 | .009 | 0.466 |
| Block X Age Group | 1 | 93 | 17.302 | .157 | **<.001\*** |
| tDCS Condition X Block X Age Group | 3 | 279 | 5.729 | .058 | **.001\*** |

For accuracy, the ANOVA showed significant main effects of Block and Age Group, and a significant interaction effect of Block * Age Group. Other main and interaction effects were not significant (Table S1.2). Post hoc tests revealed significantly higher error counts in block 6 compared to block 1 across tDCS conditions for young, but not elderly participants. Young participants made significantly more errors than elderly participants in blocks 1 and 7-8 with sham tDCS, block 6 with 1mA, blocks 3 and 5-6 with 2mA, and block 6 with 3mA tDCS (Figure S2.2).

The ANOVA for RT variability showed significant main effects of Block and Age Group. No other main and interaction effects were significant (Table S1.3). Post hoc tests revealed significantly larger RT variability in elderly than young participants in all blocks of all tDCS conditions (Figure S2.3).

### 2.2. Recall: Effects of Age and tDCS Intensity on SRTT Performance Post-Learning

At recall, the ANOVA for absolute RTs showed significant main and interaction effects of Block, Age Group, Block * Age Group, and Block * tDCS Condition. Remaining main and interaction effects were not significant (Table 1). Post hoc tests showed significantly faster RTs in recall blocks 2-3 compared to the within-condition recall block 1 for all groups and conditions. The young sample’s RTs were significantly faster than the elderly sample’s at all blocks following all tDCS conditions. Compared to sham, young participants had significantly faster RTs following 1mA tDCS at recall blocks 2-3. Elderly participants’ RTs at recall did not differ significantly between active and sham tDCS (Figure 1).

**Figure 1:**
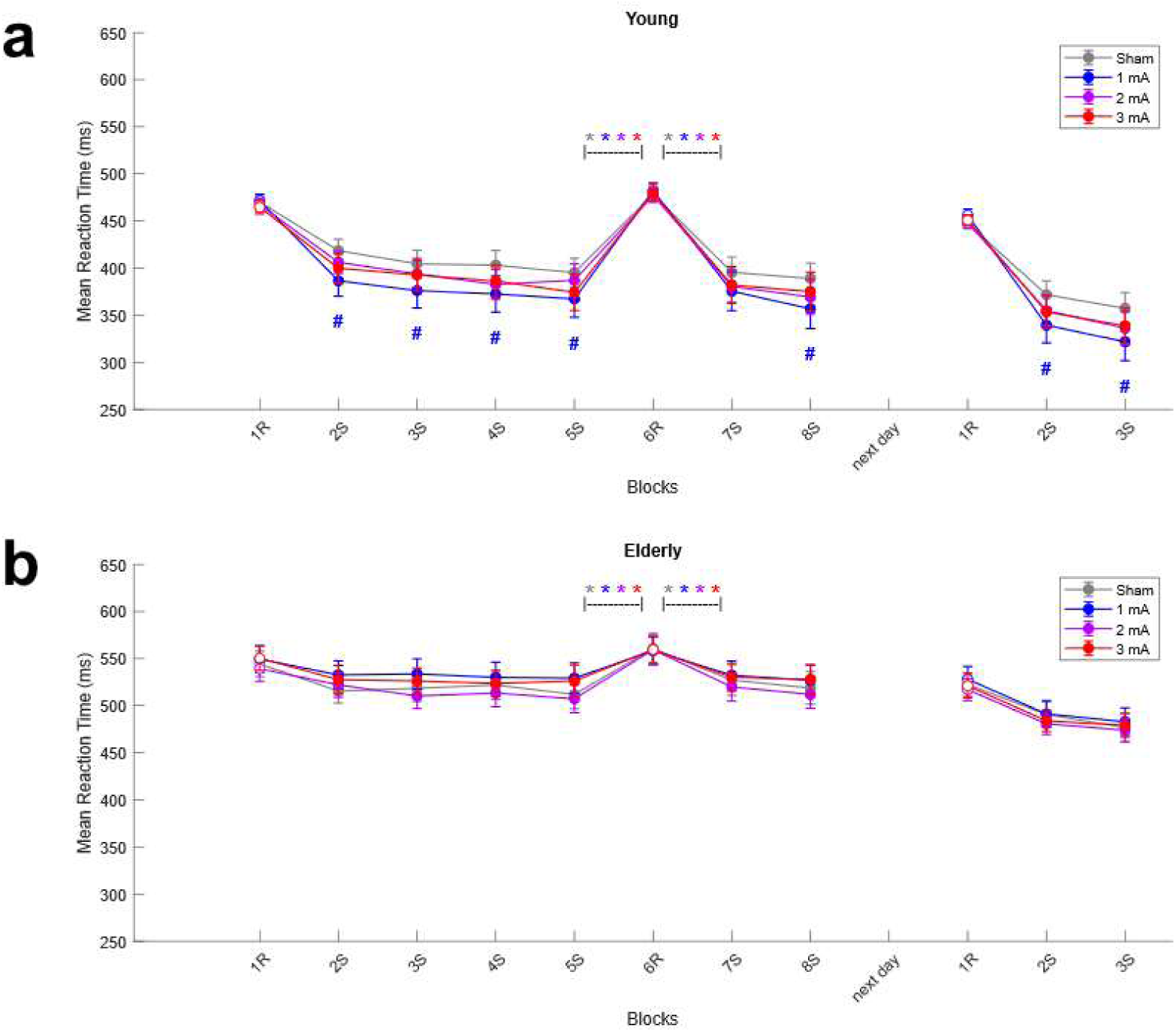
Absolute Reaction Times (RTs) of the Learning and Recall Components of the Serial Reaction Time Task (SRTT) in Young (a) and Elderly (b) participants. Online anodal-tDCS was applied during the main SRTT task (first 8 blocks). The next day, approximately 24 ± 2 hours later, participants completed 3 recall blocks of the SRTT without tDCS. The mean baseline-standardised reaction time per block with standard errors of the mean (SEM) is plotted. S = sequence blocks; R = random blocks. Paired t-test significances are indicated as: * = p<.05 between blocks 5S & 6R and blocks 6R & 7S within stimulation conditions, where grey refers to the sham tDCS condition, blue to 1mA tDCS, purple to 2mA tDCS, and red to 3mA tDCS; # = p<.05 between sham & 1mA tDCS; Filled markers indicate p<.05 compared to the respective block 1R within the main and recall SRTT, analysed separately for each age group and tDCS condition.

For baseline-standardised RTs, the ANOVA showed significant main and interaction effects of Block, Age Group, tDCS Condition, Block * Age Group, and tDCS Condition * Age Group. Remaining interaction effects were not significant (Table S1.1). For both age groups, post hoc tests revealed significantly faster baseline-standardised RTs in recall blocks 2-3 compared to the within-condition recall block 1 across tDCS conditions. Young participants had significantly faster baseline-standardised RTs than elderly participants at recall blocks 2-3 across tDCS conditions. For young adults, baseline-standardised RTs were significantly faster at recall blocks 2-3 following 1mA tDCS than sham. Older adults’ baseline-standardised RTs at recall did not differ significantly between active and sham tDCS (Figure S2.1).

The ANOVA for accuracy at recall showed significant main and interaction effects of Block and Block * Age Group (Table S1.2). Other main and interaction effects were not significant. In young participants, compared to the within-condition recall block 1, post-hoc tests revealed significantly lower error counts in recall block 2 of all tDCS conditions and recall block 3 following sham, 2mA, and 3mA tDCS. In elderly participants, error counts were significantly lower in block 2 following 2mA and 3mA tDCS compared to the within-condition recall block 1. Young participants made significantly more errors than elderly participants at recall block 1 following sham tDCS (Figure S2.2).

For RT variability at recall, the ANOVA showed significant main and interaction effects of Block, Age Group, and Block * Age Group (Table S1.3). No other main and interaction effects were significant. Post hoc tests revealed that elderly participants had significantly larger RT variability than young participants in all recall blocks following all tDCS conditions (Figure S2.3).

**Figure 2:**
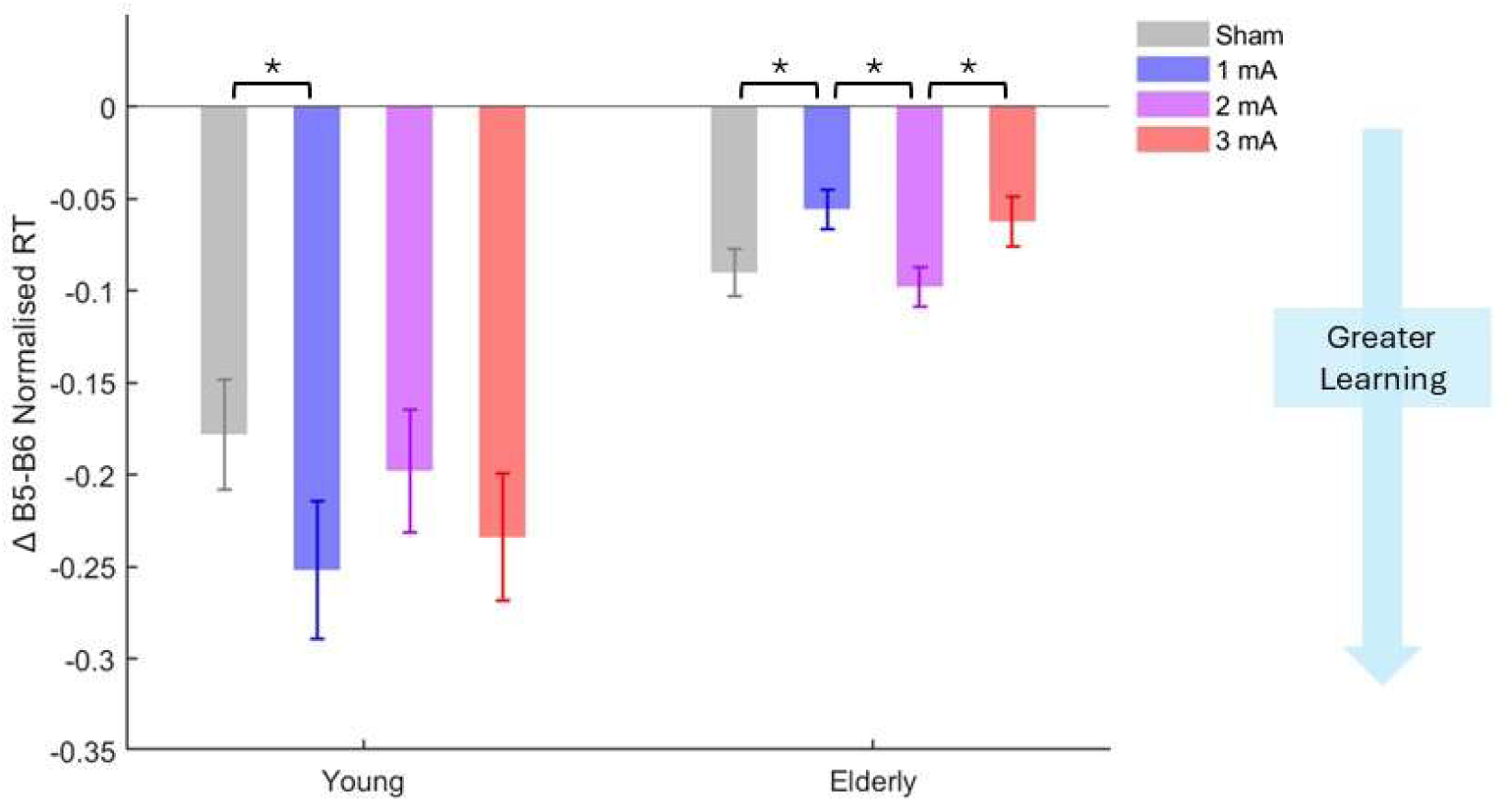
Magnitude of Selective Sequence Learning during the Learning Component of the Serial Reaction Time Task (SRTT) in Young and Elderly Participants. Anodal tDCS was applied during the SRTT. The mean learning effect (delta of sequence block 5 – random block 6 using baseline-standardised mean reaction times) with standard errors of the mean (SEM) is plotted. There was a significant main effect of age group (F(1, 93)= 20.275, p<.001), and interaction effect of tDCS condition and age group (F(2.81, 261.29)= 5.819, p=.001). Paired t-test significances are indicated as: * = p<.05 between tDCS conditions.

### 2.3. tDCS Blinding and Side-Effects

The *χ*^2^ analyses revealed that heterogeneity in guessing tDCS intensity correctly could not be assumed across age groups, with *χ*^2^(3) = 38.625, p<.001. The secondary *χ*^2^ analysis for young adults revealed the same, with *χ*^2^(3) = 15.681, p=.001 (Cross Table S2.1). Standardised residuals indicated that young participants guessed the stimulation intensity incorrectly significantly more often than expected by chance following 3 mA tDCS (z = 2.0), while guessing accuracy was heterogenous for all other tDCS conditions (−1.96 > z > 1.96). For older adults, heterogeneity could also not be assumed, *χ*^2^(3) = 34.043, p<.001 (Cross Table S2.1). Standardised residuals indicated that they correctly identified the sham condition significantly more often than expected by chance (z = 4.3), while guesses in all other tDCS conditions were heterogenous (−1.96 > z > 1.96).

Reported side-effects during and post tDCS were minor across conditions (Summary Table S2.2 & S2.3; ANOVA Table S2.3 & S2.4). For side-effect scores, ANOVAs revealed significant main and interaction effects of Age Group, tDCS Condition, and Age Group * tDCS Condition (p<.05) for ‘itching’, ‘tingling’, and ‘pain’; significant main effects of tDCS Condition and Age Group for ‘visual phenomena’, ‘burning’, and ‘redness’; significant main effects of Age Group for ‘headache’, ‘fatigue’, ‘difficulty concentrating’, and ‘nervousness’; and no significant effects for ‘sleep problems’. Post hoc tests showed that young participants scored ‘itching’, ‘tingling’, ‘burning’, ‘pain’, ‘fatigue’, and ‘difficulty concentrating’ significantly higher than elderly across tDCS conditions, as well as ‘redness’ with sham and 2mA, ‘headache’ with 2mA, ‘visual phenomena’ with 3mA, and ‘nervousness’ with sham, 1mA, and 2mA tDCS. Compared to sham, young participants scored ‘itching’, ‘tingling’, and ‘redness’ significantly higher with 2mA and 3mA tDCS, ‘visual phenomena’ with 2mA, ‘burning’ with 1mA and 3mA, and ‘pain’ with all anodal tDCS conditions. Compared to sham, elderly participants scored ‘itching’ significantly higher with 2mA and 3mA tDCS, and ‘tingling’ with 3mA.

## 3. Discussion

This study titrated anodal M1 tDCS intensities during IMSL in young and older adults. The tDCS was well tolerated with minimal side effects. Both age groups showed IMSL in all tDCS conditions, with SRTT RTs decreasing over sequence blocks while remaining significantly slower in random blocks. Young adults had consistently faster RTs than older adults (Figure S1.1) and greater selective sequence learning across tDCS conditions, assessed by comparing the difference of baseline-standardised RTs of a learnt-sequence block with those of an unlearnt random block. In young adults, 1 mA tDCS improved IMSL relative to sham, with faster RTs in the SRTT and superior selective sequence learning, and memory in the recall condition. Compared to sham, absolute RTs of older adults were unaffected by anodal tDCS, but 1mA tDCS acutely impaired selective sequence learning. At recall, both age groups showed successful IMSL consolidation across conditions, with significantly faster RTs in sequence blocks than in the random block.

### 3.1. Online 1mA Anodal tDCS facilitates Implicit Motor Sequence Learning in Young Adults

Young adults had significantly faster RTs in sequence blocks of both learning and recall SRTTs with 1mA tDCS compared to sham. Additionally, 1mA tDCS led to greater selective sequence learning compared to sham. These results align with previous literature reporting improved motor sequence learning with anodal tDCS over M1 in healthy young adults, measured by RTs during learning and consolidation [4,7,20,28–30,62]. Particularly 1mA anodal tDCS consistently enhances IMSL in young adults, as replicated here [4,63–65]. However, 2mA tDCS yielded no RT improvements, consistent with Kang & Paik (2011) who found no SRTT IMSL enhancement with 20 min anodal M1 tDCS [29]. To our knowledge, 3mA tDCS had not previously been applied to healthy young adults for IMSL, for which performance was unaffected.

### 3.2. Young Adults Outperformed Older Adults in SRTT Reaction Times and Selective Sequence Learning, but not Accuracy

Consistent with prior work, young adults had faster RTs in the SRTT than older adults (Figure S1.1) [15,66,67]. Comparable to Harrington and Haaland (1992), young adults showed greater selective sequence learning than older adults [66]. Additionally, young participants had better recall performance than older participants, with faster absolute and baseline-standardised RTs (Figures S1.1 and S1.2). Unlike Harrington and Haaland (1992), we observed significant effects of Age on SRTT accuracy [66]. During learning, accuracy was similar between age groups, but young participants made significantly more errors than elderly in the interrupting random block 6 in anodal tDCS conditions, but not sham, likely reflecting an improvement in selective sequence learning from anodal tDCS in young adults (Figure S1.3). RT variability was unaffected by tDCS, but lower in young adults, aligning with Harrington & Haaland (1992) [66].

### 3.3. Temporary Age-Dependent Impairments of Selective Sequence Learning with 1mA Anodal tDCS

In contrast to young adults, the elderly group showed no significant RT changes for general learning between anodal and sham tDCS conditions. However, their selective sequence learning was significantly reduced by 1mA tDCS compared to sham. At recall, differences between tDCS conditions in the older sample were no longer present, with unchanged baseline-standardised sequence block RTs relative to sham. Reduced tDCS effects in healthy elderly participants have previously been reported with intensities of 1-3 mA, for both motor sequence learning and cortical excitability [11,35–37,68].

### 3.4. Proposed Mechanisms of Action

Like tDCS-induced plasticity, the induction and maintenance of LTP is dependent on glutamatergic signalling [8,69–71]. Glutamatergic NNMDARs regulate intracellular Ca2+ concentrations, determining plasticity direction (LTD or LTP) [47,72–74]. The acquisition of motor memories depends on glutamatergic LTP and its strengthening of task-relevant synapses [47,75,76]. When glutamatergic activity is disrupted, LTP and motor learning are impaired [33,77–79]. Similarly, the NMDAR antagonist dextromethorphan abolishes anodal tDCS facilitation of cortical excitability [70]. The inhibitory neurotransmitter GABA gates glutamatergic plasticity and contributes to motor learning, especially early acquisition [7,80]. Motor learning is driven by changes in the E/I balance of glutamate and GABA in M1, with decreases accomplished by GABAergic inhibition and increases resulting from glutamate activity. Greater E/I ratios correlate with better motor learning [35,41,81], while pharmacologically enhancing GABAergic inhibition in humans impairs motor sequence learning [77,82].

These mechanisms may explain why anodal tDCS of 1-3mA increases cortical excitability [9], while only 1mA facilitated IMSL in our young sample. Anodal tDCS reduces GABA in target areas, disinhibiting glutamatergic LTP [8,41,83]. Lower intensities (1mA) may allow greater glutamatergic activation of task-relevant synapses. However, tDCS effects are non-focal and therefore additionally disinhibit task-irrelevant synapses [84,85]. At higher intensities, this may cause diffuse and noisy LTP. A decreased signal-to-noise ratio is inefficient for learning, as demonstrated in the rodent hippocampus, where the saturation of LTP through cross-bundle tetanisation impaired spatial learning [86]. While this tetanisation was suprathreshold, the subthreshold 2mA and 3mA anodal tDCS in this study may have induced enough synaptic noise to antagonize the IMSL benefits observed with 1mA tDCS.

These mechanisms imply glutamatergic overactivation in the effects of higher anodal tDCS intensities on IMSL. In accordance, non-linear dose-dependent effects of glutamatergic NMDAR activation on NIBS-induced neuroplasticity, LTP, and motor learning have been demonstrated before, with glutamatergic overactivation impairing plasticity induction and learning in humans [87–91]. Such glutamatergic overactivation has been linked to age-related disorders, like Alzheimer’s disease, where amyloid beta oligomers activate a signalling cascade excessively increasing calcium influx through NMDARs until apoptosis [92].

In this study, older adults showed slower RTs, reduced selective sequence learning, and greater RT variability compared to young adults. This is consistent with age-related neuroplasticity declines due to structural, molecular, and neurotransmitter-system changes [2,93–96]. Advancing age alters GABAergic and glutamatergic systems, reducing neurotransmitter levels and NMDAR function and availability [44–48,97,98]. Due to the dynamic E/I changes in learning, it is difficult to separate glutamatergic and GABAergic activity in explaining the missing tDCS effects in elderly. If solely caused by glutamate decreases, higher anodal tDCS intensities decreasing GABAergic inhibition of glutamatergic activity should have facilitated behavioural performance. This was not observed and leaves the selective sequence learning impairment in older adults under 1mA tDCS unexplained. Decreased GABA concentrations may therefore contribute to altered tDCS responses in older adults. Reduced GABA leads to hyperexcitability and impaired cognitive function in healthy and pathological aging [99,100]. King et al. (2020) showed that only older adults with high baseline GABA showed learning-related decreases in GABA comparable to young participants, associated with greater motor sequence learning. Thus, age-related decreases in GABA likely caused synaptic hyperexcitability, reducing task-relevant signal-to-noise ratios and impairing performance, reflected by reduced selective sequence learning compared to young adults. Therefore, anodal tDCS may be inefficient for improving IMSL in older adults, due to disinhibition.

In older adults, selective sequence learning was only acutely reduced during learning, not at recall. Apart from neurochemical alterations, a key factor to consider is functional network activation and connectivity, which anodal tDCS acutely alters [101]. During ageing, the connectivity of the core functional network decreases and interhemispheric connectivity increases [50,83,102]. Compared to young adults, older adults show greater activations of classical motor control regions and additional recruitment of higher-level sensorimotor and frontal regions during motor tasks [102]. Increased task-related recruitment of frontal, motor, and sensorimotor regions in the elderly is associated with better motor performance and higher baseline GABA levels [35,101,102]. This aligns with the compensation hypothesis, which argues that increased activation and additional recruitment of cerebral areas allow elderly to maintain cognitive performance comparable to young adults, despite age-related functional and physiological deficits [102–104]. Therefore, older adults may require additional frontal and sensorimotor targets for tDCS-induced motor learning facilitation.

### 3.5. Limitations and Future Directions

Several limitations of the present study require acknowledgement. Chi-square analyses indicated ineffective blinding for elderly in the sham tDCS condition, and young participants guessed the wrong intensity following 3mA tDCS more frequently than chance. However, elderly participants’ performance under sham differed significantly from 1mA tDCS during learning, in contrast to all active tDCS conditions during learning and recall, arguing against a systematic effect of unblinding. In terms of side effects, only itching and tingling scores differed significantly between sham and active tDCS, with all mean scores <1 of 5, indicating very mild sensations. Additionally, although young participants reported greater side effects in the 3mA condition, they guessed lower intensities, possibly undermining the effects of 3mA tDCS, for which performance remained comparable to sham. Altogether, despite topical anaesthesia, somatosensory and expectation effects cannot be fully ruled out. Due to significant differences between age group and tDCS condition side effect scores, we analysed a linear mixed model for the impact of side effect scores on selective sequence learning, controlling for session number (Table S2.5). Here, side effects had no significant effect on selective sequence learning for either age group, suggesting they did not significantly alter performance.

Regarding the SRTT, there is no clear consensus on monitoring individuals’ awareness to learning, specifically to exclude explicit learning [55,105]. While formerly believed that implicit learning recruits striatal regions and explicit learning competitively the medial temporal lobe, both forms of learning rely on hippocampal activity and related structures for higher-order associative learning [106,107]. Nevertheless, sequence awareness does not shift activation towards the medial temporal lobe, nor does it significantly affect motor sequence learning performance in young adults [106,107]. This suggests that sequence awareness does not primarily drive tDCS modulation of motor sequence learning in young adults. In older adults, however, explicit sequence awareness might impair IMSL, making it a relevant factor to consider in our results [108].

Additionally, although underlying neurophysiology provides a mechanistic interpretation of our results, direct neurophysiological measures were not conducted. Future research would benefit from further exploring tDCS optimisation in aging, using direct neurophysiological paradigms such as pharmacological interventions, neuroimaging, and additional forms of non-invasive brain stimulation. One such example could involve titrating tDCS intensities and using magnetic resonance spectroscopy to quantify glutamate and GABA in both young and older adults during IMSL to understand their contributions to behavioural tDCS effects more clearly.

## 4. Conclusion

We demonstrated that anodal tDCS intensity has non-linear behavioural effects on motor sequence learning that are relevantly shaped by age. Online 1mA stimulation selectively facilitated motor performance, sequence learning, and memory in young adults, while acutely impairing selective sequence learning in older adults. These divergent effects are best understood within age-related alterations of the E/I balance of glutamate and GABA constraining the behavioural efficacy of tDCS. The findings underscore the necessity for population-optimised stimulation protocols for experimental research and clinical tDCS applications, especially regarding age-related disorders that alter E/I neurotransmission, such as Alzheimer’s Disease.

## 5. Methods and Materials

We conducted a randomised, double-blind, sham-controlled, crossover study. Participants completed four counterbalanced experimental sessions corresponding to different tDCS conditions. Procedures were in accordance with the statements of the Declaration of Helsinki and ethical approval was granted by the ethics committees of the Leibniz Research Centre for Working Environment and Human Factors, Dortmund, Germany (Approval 09.10.2018, Code: 151, Amendment 14.07.2020) and the Faculteit Geneeskunde en Levenswetenschappen, Hasselt University, Belgium (Approval 26.03.2019, Code: B9115201939823).

### 5.1. Sample

The study included 96 healthy participants; 48 were elderly (26 females; mean age 71.15 ± SD 5.12 years) and 48 young adults (26 females; mean age 25.19 ± SD 3.68). Sample size was determined using a power analysis (G*Power v.3.1.9.7) [51], based on a repeated measures ANOVA for the interaction effect between Age Group (between-subject) and tDCS Condition (within-subject). Assuming Cohen’s f = 0.25 (equivalent to *η*^2^*p*= 0.06), alpha = 0.05, and power = 0.95, the required sample size was N = 36 (18 per age group). This was increased to 48 participants per group to counterbalance tDCS conditions and SRTT-sequence order.

All participants were right-handed non-smokers. Handedness was assessed with the Edinburgh Handedness Inventory [52]. Medical screening by a physician excluded tDCS contraindications: History of neurological or psychiatric disorders, epilepsy, seizures, CNS-acting medications, metal implants in head and neck areas, substance-abuse, pregnancy, breastfeeding, skin disease or injury in stimulation areas, or any severe internal medical disorder. For elderly participants, cognitive deficits were ruled out using the Montreal Cognitive Assessment (MoCA) [53]. All participants provided written informed consent and received financial compensation.

Of the elderly sample, the data of N = 23 were previously collected and published by our group [37] using the same inclusion criteria, materials, and tDCS-SRTT procedure. The remainder of this study’s participants were newly recruited.

### 5.2. Transcranial Direct Current Stimulation

Transcranial direct current stimulation was delivered by a constant current stimulator (NeuroConn, Germany) using conductive rubber electrodes (5×7 cm, 35 cm^2^) with Ten20 Conductive Paste (Weaver, USA) applied to the scalp to minimise impedance and maintain electrode position. The anode was placed over the left M1 hand area (C3 according to the international 10-20 EEG system) with the long electrode edge oriented 45° to the midline. C3 was identified by first marking the vertex (Cz) at the intersection of nasion–inion and left–right preauricular distances, followed by marking C3 at 20% of the preauricular distance from Cz toward the left preauricular point. The cathode was placed over the contralateral supraorbital area.

Before tDCS, a topical anaesthetic cream was applied to stimulation sites to blind participants by reducing somatosensory sensations during stimulation [54]. EMLA® cream (2.5% lidocaine and 2.5% prilocaine; Aspen Pharma Trading Limited, Dublin, Ireland) or lidocaine HCL 2.5% buffered cetomacrogol cream (Aa.Pharma Oud-Turnhout, Hasselt, Belgium) was used for both groups.

The stimulation was controlled by an external trigger (flip-flop mode). All stimulation conditions had 30-seconds ramp-up and 30-seconds ramp-down. For active tDCS conditions, a current of 1mA, 2mA, or 3mA was applied for the entire SRTT duration. Sham tDCS included 30-seconds ramp-up, then 1 mA current for 30 seconds, followed by 30-seconds ramp-down and no stimulation for the remaining session [5]. To prevent carry-over effects, sessions were separated by at least 7 days.

### 5.3. Serial Reaction Time Task for Implicit Motor Sequence Learning

Participants completed the SRTT in E-Prime 2.0 (Psychology Software Tools Inc., Sharpsburg, USA). Four horizontal dashes appeared on screen, with a circle marking one of the four positions (Figure 3B). Using their right hand, participants pressed the corresponding key on a 4-key response box according to the circle’s position (keys 1-4 for index, middle, ring, and little finger respectively). Participants were instructed to perform the task as quickly and accurately as possible. The next stimulus was only shown 500ms after responding, regardless of accuracy. No feedback was given.

**Figure 3.**
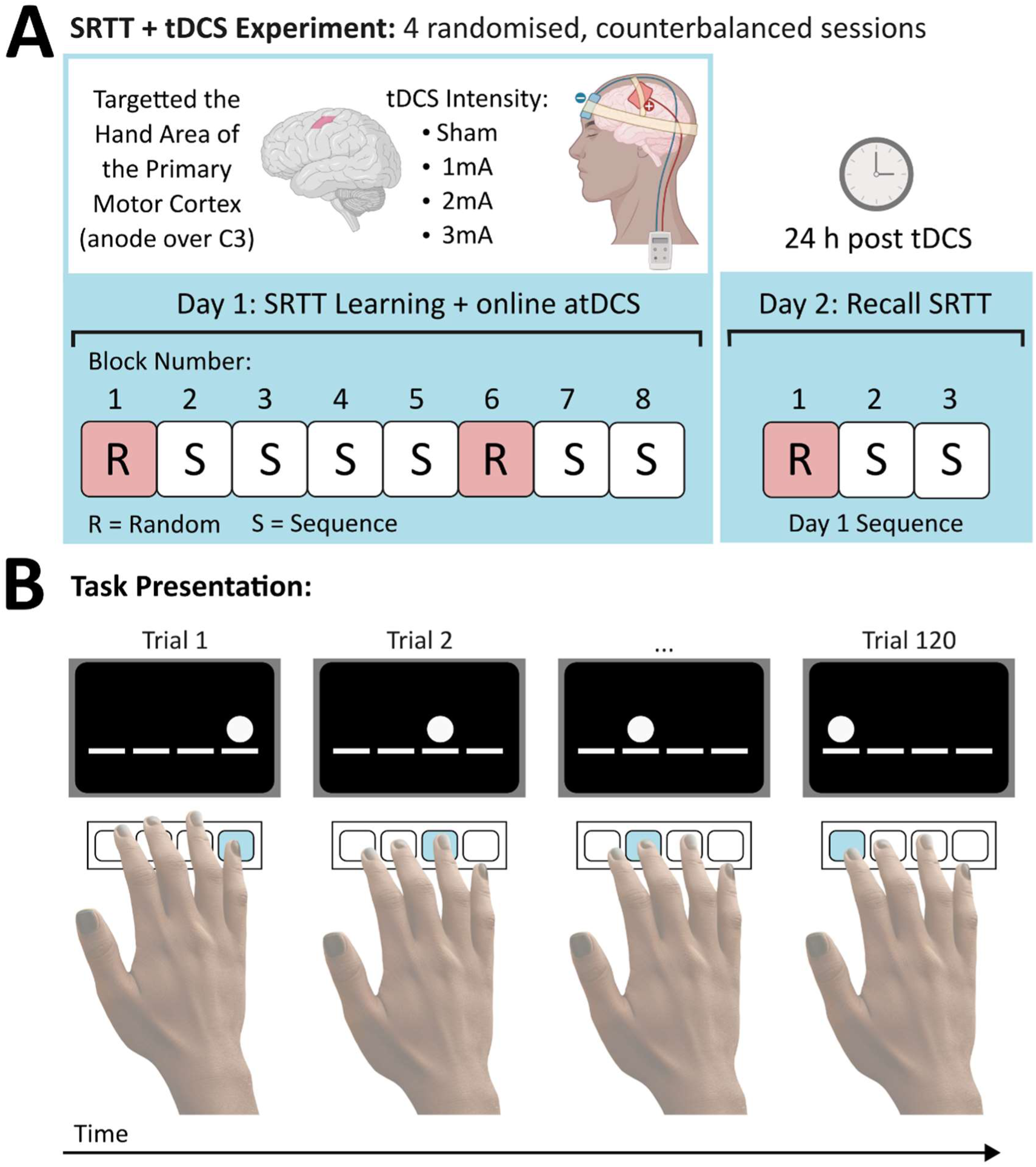
Experimental Course. A: All participants took part in 4 randomised, counterbalanced sessions. During the main serial reaction time task (SRTT), anodal tDCS was applied over the hand area of the primary motor cortex (anode over C3 in the 10-20 EEG system and cathode over the contralateral supraorbital area) at intensities of 1mA, 2mA, 3mA, and sham. The main SRTT included 8 blocks, of which blocks 1 & 6 included 120 pseudorandomised items without a sequence and the remaining blocks a repeating 12-item sequence. Blocks are labelled as R = random and S = sequence block. Each session contained a new counterbalanced sequence. The next day, three SRTT recall blocks were conducted. Recall-block 1 was a new pseudorandom 120-item block and recall-blocks 2 & 3 had the previous day’s repeating 12-item sequence. **B:** The task was presented on a computer monitor. Each finger was associated with a response key (keys 1-4 for index, middle, ring, and little finger respectively), and each response key with a visual cue consisting of 4 lines. When a circle appeared at one of the line positions, the participant was tasked with pressing the corresponding key as quickly and accurately as possible (Figure created with Inkscape and Biorender.com).

The experiment was carried out across two consecutive days. Day 1 included the 8-block SRTT learning with online tDCS. Blocks 1 and 6 included stimuli in random order. Blocks 2–5 and 7–8 were sequence blocks to induce IMSL [55]. Sequences consisted of a fixed 12-item sequence repeated 10 times per block. Participants were not told about the repeating sequence. To avoid learning between tDCS conditions, the experiment included four unique sequences, counterbalanced and randomised between stimulation sessions. Random blocks included 120 stimuli in pseudorandomised order.

The second day, participants returned for a condensed 3-block SRTT (recall). This included one random block followed by two blocks with the same sequence used the previous day. No tDCS was applied during recall.

### 5.4. Experimental Procedure

Participants sat in front of a 19” monitor (Samsung Syncmaster 957p, South Korea) with 50 cm eye-distance. They first completed a pseudorandom 20-item SRTT practice block to become familiar with the task. Once completed, they began the main SRTT (8 blocks). The SRTT was synchronized with tDCS onset, such that the first trial began following the ramp-up. Stimulation ramped down after the final trial. To assess blinding and side-effects, participants completed a questionnaire rating 10 side effect items during and after stimulation on a scale of 0-5 (none to extremely strong) and were asked to guess which tDCS intensity they received on an ordinal scale of 0-3 mA [56,57]. The next day, 24 ± 2 hours post tDCS administration, participants returned for the recall SRTT (Figure 3; experimental course). Following completion of all sessions, participants reported whether they noticed a sequence in any session.

### 5.5. Statistical Analyses

Data were pre-processed in Matlab (v9.9, R2020b, The MathWorks Inc., M.A., USA) and statistically analysed using SPSS (v25, IBM Corp., Armonk, N.Y., USA) and R Statistical Software (v4.4.2; R Core Team 2024).

For each participant and tDCS condition, mean and standard deviations (SD) of correct trial reaction times (RT) were calculated per block. Outliers >3 SDs from the individual mean were excluded. Mean reaction times (mRT) per block were standardised onto baseline block 1 (mRT _Block x_ / mRT _Block 1_) for each participant and tDCS condition. The mean error count per block was calculated to reflect accuracy. Participants with <50% accuracy were excluded from analyses. RT variability was quantified as the SD of correct RTs per block, post outlier removal.

RTs and error counts were the main SRTT outcome variables. RTs, baseline-standardised RTs, RT variability, and error counts were analysed in separate mixed repeated measures ANOVAs with within-subject factors block (8 levels for SRTT learning, 3 levels for recall) and tDCS condition (4 levels: sham, 1 mA, 2 mA, 3 mA), and the between-subject factor age group (2 levels: young and elderly). For all ANOVAs, the assumption of sphericity was checked using the Mauchly test, and when necessary, Greenhouse-Geisser corrections were applied. In case of significant effects, ANOVAs were followed up by exploratory, uncorrected post-hoc *t*-tests (two-tailed, p < 0.05).

Both unspecific motor learning (key presses) and sequence learning (order of key presses) decrease SRTT RTs. As compared to RTs in sequence blocks (here 2-5), these increase abruptly in an interrupting random block (here block 6), with the difference selectively reflecting sequence learning, rather than unspecific motor learning, as the motor component of the task remains unchanged [58–61]. Therefore, blocks 5 vs 6 are of major interest for selective sequence learning, comparing sequence learning-dependently decreased RT vs RT independent of sequence learning. We conducted a secondary mixed ANOVA with factors Block (2 levels: blocks 5 & 6), tDCS Condition (4 levels), and Age Group (2 levels) on baseline-standardised RTs to compare selective sequence learning. Only baseline-standardised RTs were analysed, to account for age-related differences in baseline RT. Post hoc comparisons examined the RT difference of block 5 vs 6 per participant as an index of selective sequence learning (Equation 1).

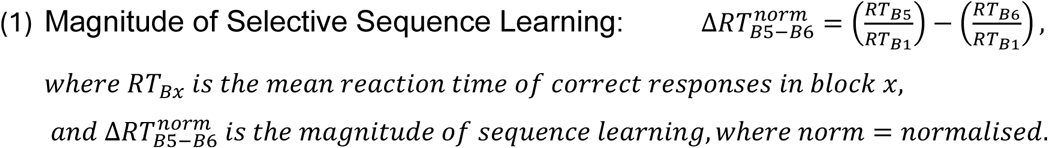

Blinding was assessed by a Chi-square test on the binary outcome of whether participants guessed tDCS intensity correctly or not. Follow-up Chi-square tests were conducted separately by age group to examine age-specific patterns, with post hoc standardised residuals derived from age-specific contingency tables. For analysing tDCS side-effects, scores for each item were analysed separately using a repeated measures ANOVA with within-subject factor tDCS condition (4 levels) and between-subject factor age group (2 levels).

## 6. Data Availability

The datasets generated during and/or analysed during the current study are not publicly available due to ethical restrictions and to ensure participant privacy but are available from the corresponding author on reasonable request.

## 9. Acknowledgements

We thank the participants who took part in this experiment for their help in advancing scientific research. We also extend our gratitude to our colleagues who assisted through technical support: Daniel Strobel, Andreas Volgmann, Tobias Blanke, and Mark Geraerts all made the experimental setup possible. Additionally, our thanks go to Nicole Rück for helping with participant recruitment, and Lisa Göbel for her impeccable administrative structures. We are also thankful to Klaus Golka and Jan G. Hengstler who helped to medically screen participants.

## 10. Authorship Contribution Statement

**A.M.F.:** Investigation, data curation, formal analysis, visualisation, writing – original draft, writing - review and editing; **R.U.:** Investigation, writing – original draft; **E.G.-S.:** Investigation, writing – review and editing; **L.D.M.:** Investigation, writing – review and editing; **Y.X.:** Investigation; **M.B.:** Investigation; **M-F.K.:** Conceptualisation, methodology, supervision, writing – review and editing; **R.L.J.M.:** Conceptualisation, funding acquisition, resources, supervision, writing – review and editing; **M.A.N.:** Conceptualisation, methodology, funding acquisition, resources, supervision, writing – review and editing. All authors have read and agreed to the published version of the manuscript.

