## Supplementary Figures and Tables for "Age-dependent Effects of Titrating Anodal Transcranial Direct Current Stimulation (tDCS) Intensity on Motor Sequence Learning"

### Supplementary Documents

### 1. Results of SRTT Learning and Recall

**Table 1.1: Results of the Mixed Model ANOVAs for Baseline-Standardised Reaction Time in the Serial Reaction Time Task (SRTT).**

| *Variables* | *d.f.* | *d.f.$_{error}$* | *F-value* | $\eta_p^2$ | *p-value* |
|---|---|---|---|---|---|
| **Learning (during tDCS)** | | | | | |
| tDCS Condition | 3 | 279 | 1.113 | .012 | .344 |
| Block | 1.497 | 139.205 | 60.660 | .395 | **<.001*** |
| Age Group | 1 | 93 | 20.364 | .180 | **<.001*** |
| tDCS Condition X Block | 9.766 | 908.274 | .791 | .008 | .635 |
| tDCS Condition X Age Group | 3 | 279 | 2.320 | .024 | .076 |
| Block X Age Group | 1.497 | 139.205 | 18.725 | .168 | **<.001*** |
| tDCS Condition X Block X Age Group | 9.766 | 908.274 | 1.995 | .021 | **.032*** |
| **Recall (next day)** | | | | | |
| tDCS Condition | 2.752 | 255.898 | 2.864 | .030 | **.042*** |
| Block | 1.087 | 101.064 | 88.660 | .488 | **<.001*** |
| Age Group | 1 | 93 | 23.560 | .202 | **<.001*** |
| tDCS Condition X Block | 4.161 | 387.014 | 2.273 | .024 | .058 |
| tDCS Condition X Age Group | 2.752 | 255.898 | 2.826 | .029 | **.044*** |
| Block X Age Group | 1.087 | 101.064 | 22.572 | .195 | **<.001*** |
| tDCS Condition X Block X Age Group | 4.161 | 387.014 | 2.315 | .024 | .054 |

Mauchly's test of sphericity was conducted for each ANOVA, and Greenhouse–Geisser corrections were applied where sphericity assumptions were not met. Asterisks indicate significant effects ($p<.05$). d.f. = degrees of freedom; η2$p$ = partial eta squared.

**Table 1.2: Results of the Mixed Model ANOVAs for Error Counts in the Serial Reaction Time Task (SRTT).**

| *Variables* | *d.f.* | *d.f.$_{error}$* | *F-value* | $\eta_p^2$ | *p-value* |
|---|---|---|---|---|---|
| **Learning (during tDCS)** | | | | | |
| tDCS Condition | 3 | 279 | .450 | .005 | .717 |
| Block | 4.709 | 437.933 | 10.434 | .101 | **<.001*** |
| Age Group | 1 | 93 | 4.348 | .045 | **.040*** |
| tDCS Condition X Block | 11.563 | 1075.397 | .984 | .010 | .460 |
| tDCS Condition X Age Group | 3 | 279 | .116 | .001 | .951 |
| Block X Age Group | 4.709 | 437.933 | 6.755 | .068 | **<.001*** |
| tDCS Condition X Block X Age Group | 11.563 | 1075.397 | 1.017 | .011 | .429 |
| **Recall (next day)** | | | | | |
| tDCS Condition | 2.395 | 222.745 | .727 | .008 | .508 |
| Block | 1.740 | 161.781 | 26.420 | .221 | **<.001*** |
| Age Group | 1 | 93 | .695 | .007 | .407 |
| tDCS Condition X Block | 11.563 | 1075.397 | .984 | .011 | .480 |
| tDCS Condition X Age Group | 2.882 | 268.067 | .116 | .003 | .951 |
| Block X Age Group | 4.709 | 437.933 | 6.755 | .070 | **<.001*** |
| tDCS Condition X Block X Age Group | 11.563 | 1075.397 | 1.017 | .014 | .429 |

Mauchly’s test of sphericity was conducted for each ANOVA, and Greenhouse–Geisser corrections were applied where sphericity assumptions were not met. Asterisks indicate significant effects ($p<.05$). d.f. = degrees of freedom; η2$p$ = partial eta squared.

**Table 1.3: Results of the Mixed Model ANOVAs of Reaction Time Variability (Standard Deviation) in the Serial Reaction Time Task (SRTT).**

| *Variables* | *d.f.* | *d.f.$_{error}$* | *F-value* | $\eta_p^2$ | *p-value* |
|---|---|---|---|---|---|
| **Learning (during tDCS)** | | | | | |
| tDCS Condition | 2.727 | 253.569 | .998 | .011 | .394 |
| Block | 3.048 | 283.434 | 14.063 | .131 | **<.001*** |
| Age Group | 1 | 93 | 31.192 | .251 | **<.001*** |
| tDCS Condition X Block | 7.681 | 714.373 | 1.158 | .012 | .323 |
| tDCS Condition X Age Group | 2.727 | 253.569 | .980 | .010 | .397 |
| Block X Age Group | 3.048 | 283.434 | 1.322 | .014 | .267 |
| tDCS Condition X Block X Age Group | 7.681 | 714.373 | 1.050 | .011 | .396 |
| **Recall (next day)** | | | | | |
| tDCS Condition | 3 | 279 | 1.506 | .016 | .213 |
| Block | 1.768 | 164.453 | 23.122 | .199 | **<.001*** |
| Age Group | 1 | 93 | 43.028 | .316 | **<.001*** |
| tDCS Condition X Block | 5.033 | 468.033 | .858 | .009 | .510 |
| tDCS Condition X Age Group | 3 | 279 | .356 | .004 | .785 |
| Block X Age Group | 1.768 | 164.453 | 9.271 | .091 | **<.001*** |
| tDCS Condition X Block X Age Group | 5.033 | 468.033 | 1.402 | .015 | .222 |

Mauchly's test of sphericity was conducted for each ANOVA, and Greenhouse–Geisser corrections were applied where sphericity assumptions were not met. Asterisks indicate significant effects ($p<.05$). d.f. = degrees of freedom; $\eta 2p$ = partial eta squared.

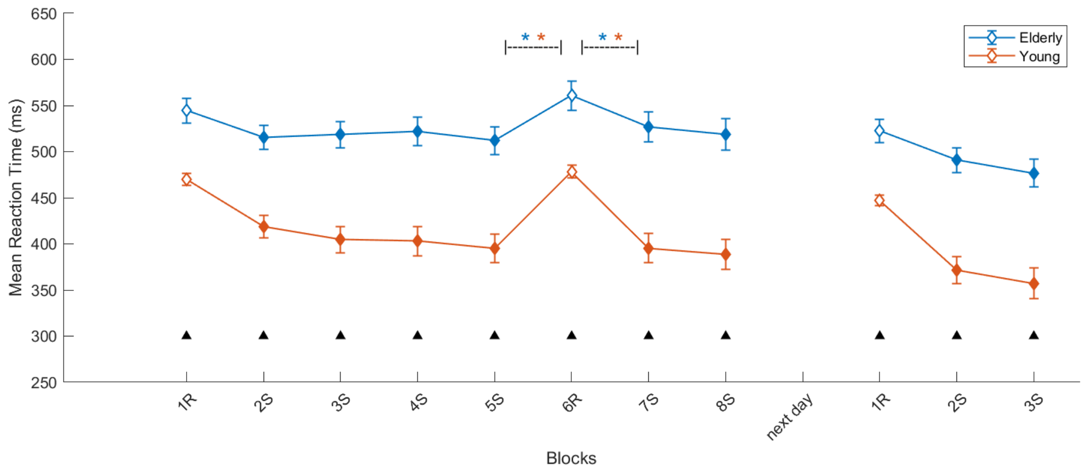

***Figure 1.1: Absolute Reaction Times (RT) in the Sham Condition of the Learning and Recall Components of the Serial Reaction Time Task (SRTT) in Young and Elderly Participants.*** Online sham-tDCS was applied during the main SRTT task (first 8 blocks). The next day, approximately 24 ± 2 hours later, participants completed 3 recall blocks of the SRTT without tDCS. The mean reaction time per block with standard errors of the mean (SEM) is plotted. S = sequence blocks; R = random blocks. Paired t-test significances are indicated as: * = p<.05 between blocks 5S & 6R and blocks 6R & 7S, where blue refers to the elderly sample, and orange refers to the young sample; ▲ = p<.05 between young and elderly participants. Filled symbols indicate p<.05 compared to the respective block 1R within the main and recall SRTT, analysed separately for each age group.

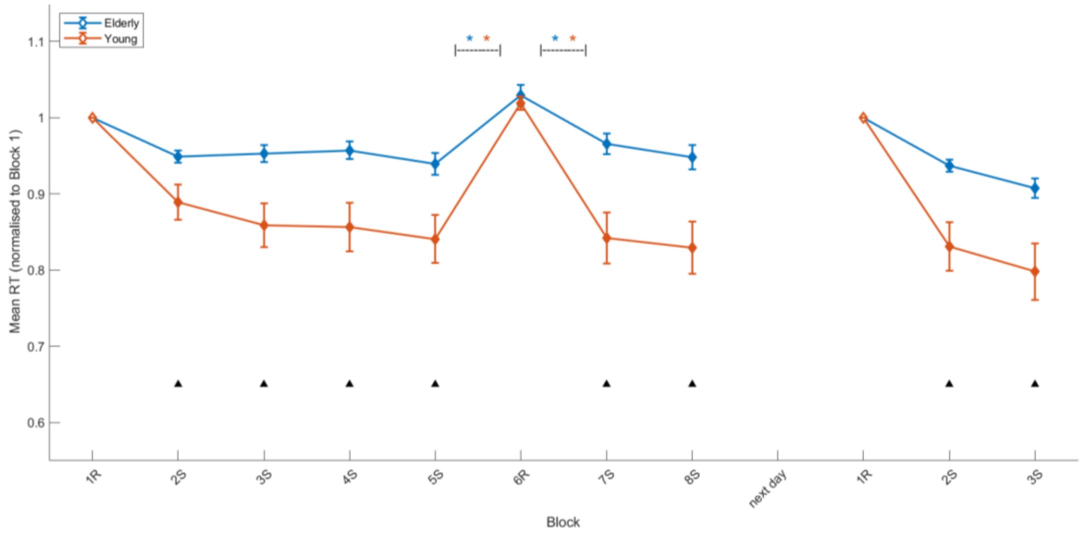

***Figure 1.2: Baseline-standardised Reaction Times (RTs) for the Sham Condition of the Learning and Recall Component of the Serial Reaction Time Task (SRTT) in Young and Elderly Participants.*** Online sham-tDCS was applied during the main SRTT task (first 8 blocks). The next day, approximately 24 ± 2 hours later, participants completed 3 recall blocks of the SRTT without tDCS. The mean baseline-standardised reaction time per block with standard errors of the mean (SEM) is plotted. S = sequence blocks; R = random blocks. Paired t-test significances are indicated as: * = p<.05 between blocks 5S & 6R and blocks 6R &7S, where blue refers to the elderly sample, and orange refers to the young sample; ▲ = p<.05 between young and elderly participants. Filled markers indicate p<.05 compared to the respective block 1R within the main and recall SRTT, analysed separately for each age group.

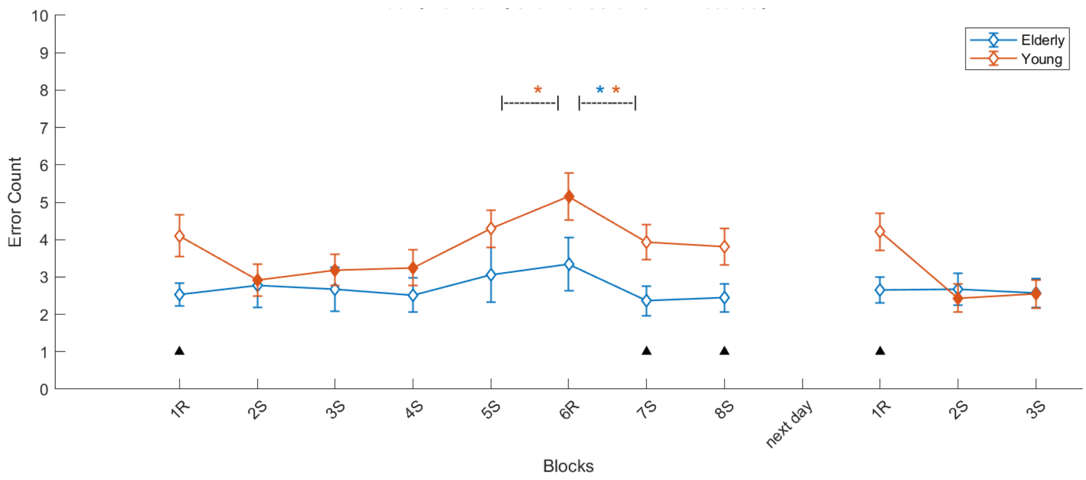

***Figure 1.3: Error Rates in the Sham Condition of the Learning and Recall Components of the Serial Reaction Time Task (SRTT) in Young and Elderly participants.*** Online sham-tDCS was applied during the main SRTT task (first 8 blocks). The next day, approximately 24 ± 2 hours later, participants completed 3 recall blocks of the SRTT without tDCS. The mean error count per block with standard errors of the mean (SEM) is plotted. S = sequence blocks; R = random blocks. Paired t-test significances are indicated as: * = $p<.05$ between blocks 5S & 6R and blocks 6R & 7S, where blue refers to the elderly sample, and orange refers to the young sample; ▲ = $p<.05$ between young and elderly. Filled markers indicate $p<.05$ compared to the respective block 1R within the main and recall SRTT, analysed separately for each age group.

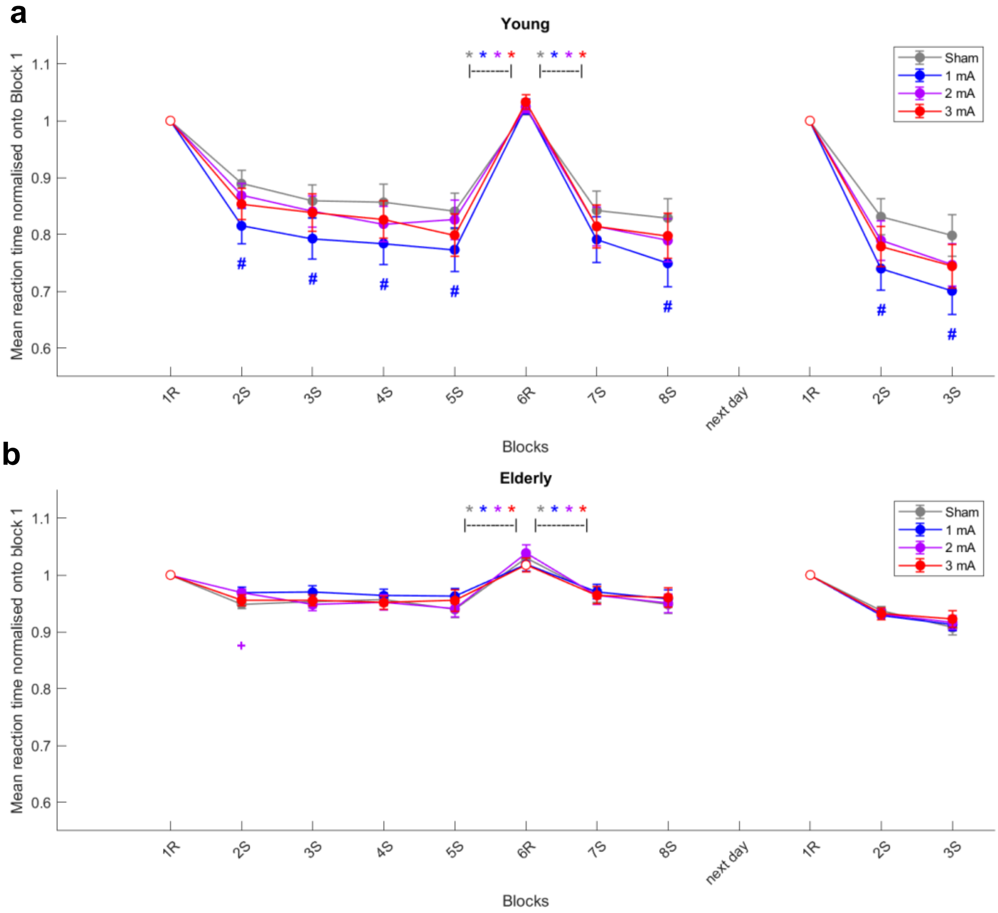

***Figure 2.1: Baseline-standardised Reaction times (RT) of the Learning and Recall Components of the Serial Reaction Time Task (SRTT) in Young (a) and Elderly (b) Participants.*** Online anodal-tDCS was applied during the main SRTT task (first 8 blocks). The next day, approximately 24 ± 2 hours later, participants completed 3 recall blocks of the SRTT without tDCS. The mean baseline-standardised reaction time per block with standard errors of the mean (SEM) is plotted. S = sequence blocks; R = random blocks. Paired t-test significances are indicated as: ***** = p<.05 between blocks 5S & 6R and blocks 6R & 7S within stimulation conditions, where grey refers to the sham tDCS condition, blue to 1mA tDCS, purple to 2mA tDCS, and red to 3mA tDCS; **#** = p<.05 between sham & 1mA tDCS; **+** = p<.05 between sham & 2mA tDCS; Filled markers indicate p<.05 compared to the respective block 1R within the main and recall SRTT, analysed separately for each age group and tDCS condition.

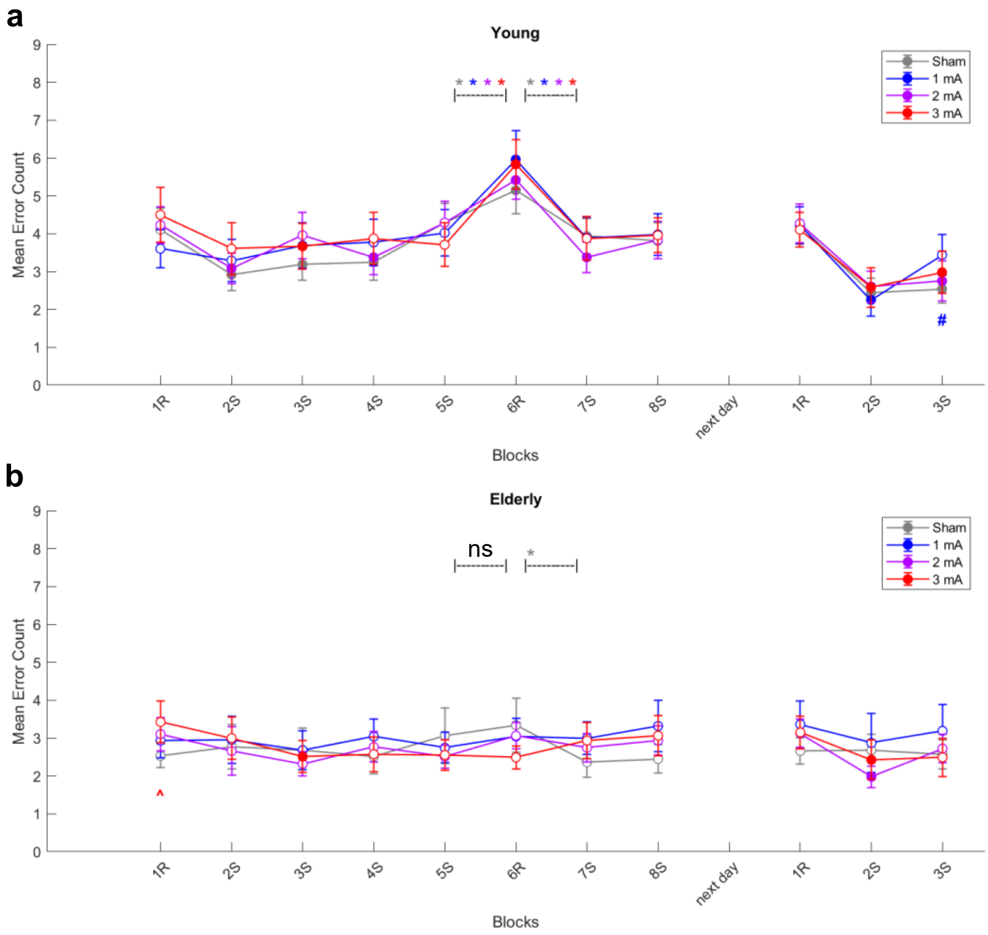

***Figure 2.2: Accuracy during the Learning and Recall Components of the Serial Reaction Time Task (SRTT) in Young (a) and Elderly (b) Participants.*** Online anodal-tDCS was applied during the main SRTT task (first 8 blocks). The next day, approximately 24 ± 2 hours later, participants completed 3 recall blocks of the SRTT without tDCS. The mean error count per block with standard errors of the mean (SEM) is plotted. S = sequence blocks; R = random blocks. Paired t-test significances are indicated as: ***** = p<.05 between blocks 5S & 6R and blocks 6R & 7S within tDCS conditions, where grey refers to the sham tDCS condition, blue to 1mA tDCS, purple to 2mA tDCS, and red to 3mA tDCS;  ns = no significant difference between blocks 5S & 6R and blocks 6R & 7S within tDCS conditions; **#** = p<.05 between sham & 1mA tDCS; **^** =  p<.05 between sham & 3mA tDCS; Filled markers indicate p<.05 compared to the respective baseline block 1R within the main and recall SRTT, analysed separately for each age group and tDCS condition.

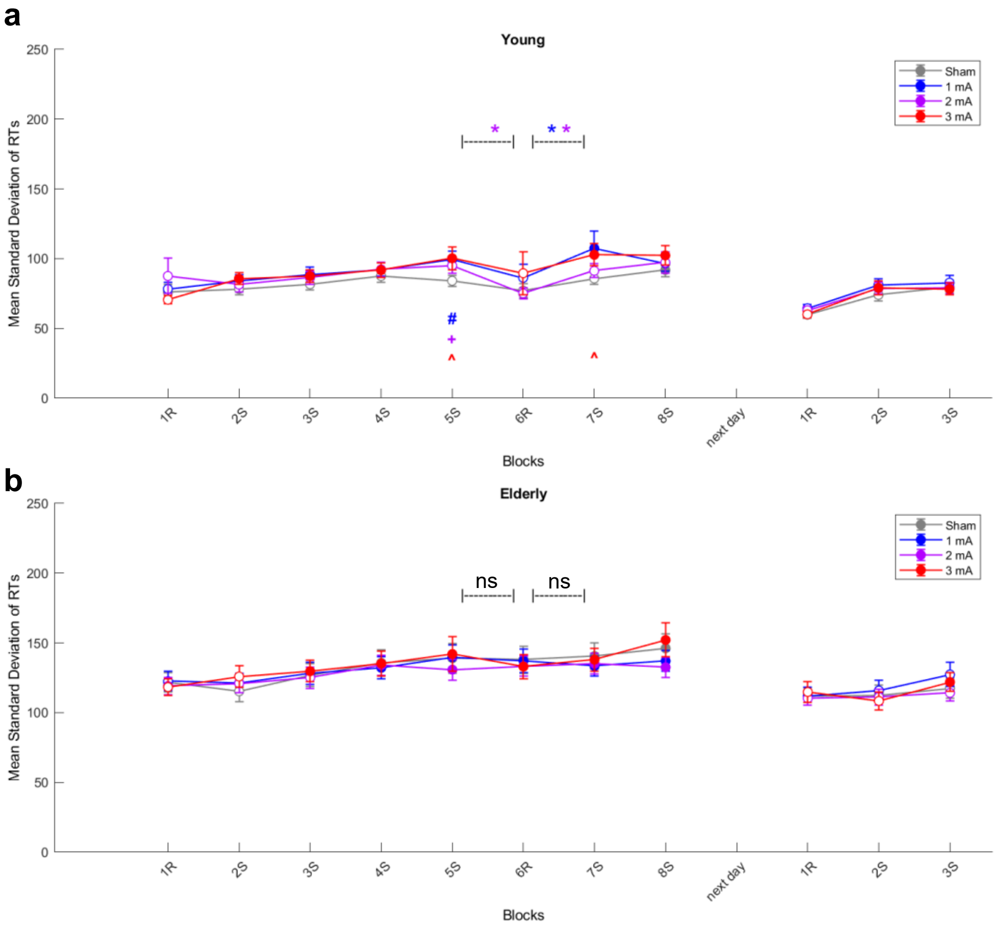

***Figure 2.3: Reaction time (RT) Variability of the Learning and Recall Components of the Serial Reaction Time Task (SRTT) in Young (a) and Elderly (b) Participants.*** Online anodal-tDCS was applied during the main SRTT task (first 8 blocks). The next day, approximately 24 ± 2 hours later, participants completed 3 recall blocks of the SRTT without tDCS. The mean standard deviation (SD) before outlier removal, excluding the first trial of each block, is plotted with standard errors of the mean (SEM). S = sequence blocks; R = random blocks. Exploratory post-hoc comparison significances are indicated as: ***** = p<.05 between blocks 5S & 6R and blocks 6R & 7S within tDCS conditions, where blue refers to the 1mA tDCS condition and red to 3mA tDCS; ns = no significant difference between blocks 5S & 6R and blocks 6R & 7S within tDCS conditions; **#** = p<.05 between sham & 1mA tDCS; **+** = p<.05 between sham & 2mA tDCS; **^** = p<.05 between sham & 3mA tDCS. Filled markers indicate p<.05 compared to the respective baseline block 1R within the main and recall SRTT, analysed separately for each age group and tDCS condition.

#### 2. tDCS Blinding and Side-Effects

**Table 2.1: Data Points of Guessed versus Received tDCS Intensity.**

| **Young Participants (N = 48)** | | | | | |
|---|---|---|---|---|---|
| | | Real tDCS Intensity | | | |
| | | Sham | 1 mA | 2 mA | 3 mA |
| Guessed tDCS Intensity | Sham | 26 | 14 | 3 | 5 |
| | 1 mA | 18 | 20 | 17 | 15 |
| | 2 mA | 4 | 11 | 22 | 20 |
| | 3 mA | 0 | 3 | 6 | 8 |
| N Correct | | **26** | **20** | **22** | **8** |
| **Elderly Participants (N = 48)** | | | | | |
| | | Real tDCS Intensity | | | |
| | | Sham | 1 mA | 2 mA | 3 mA |
| Guessed tDCS Intensity | Sham | 28 | 30 | 23 | 21 |
| | 1 mA | 10 | 10 | 11 | 16 |
| | 2 mA | 3 | 5 | 7 | 5 |
| | 3 mA | 7 | 3 | 7 | 6 |
| N Correct | | **28** | **10** | **7** | **6** |

Following each SRTT learning session, participants completed a questionnaire asking them to guess the received tDCS intensity (0, 1, 2, 3 mA). Per age group, the table compares the real tDCS intensity (columns) with the perceived tDCS intensity (rows) and shows the number of participants who guessed the respective intensity per tDCS condition, as well as the total number of correct guesses (N Correct).

**Table 2.2: Descriptive Statistics of Side-Effects during and after tDCS.**

| Side Effects | Sham | | 1mA | | 2mA | | 3mA | |
|---|---|---|---|---|---|---|---|---|
| | Young | Elderly | Young | Elderly | Young | Elderly | Young | Elderly |
| Visual Phenomena | 0.50 ± 1.05 | 0.40 ± 1.09 | 0.56 ± 1.07 | 0.23 ± 0.93 | 1.00 ± 1.54 | 0.46 ± 1.32 | 0.58 ± 1.11 | 0.10 ± 0.37 |
| Itching | 0.71 ± 1.05 | 0.00 ± 0.00 | 1.00 ± 1.24 | 0.10 ± 0.47 | 1.54 ± 1.52 | 0.17 ± 0.56 | 1.77 ± 1.49 | 0.25 ± 0.64 |
| Tingling | 0.63 ± 0.73 | 0.13 ± 0.39 | 0.92 ± 1.18 | 0.02 ± 0.14 | 1.71 ± 1.46 | 0.25 ± 0.76 | 1.50 ± 1.26 | 0.38 ± 0.91 |
| Burning | 0.52 ± 1.07 | 0.08 ± 0.35 | 0.92 ± 1.40 | 0.15 ± 0.58 | 0.90 ± 1.19 | 0.21 ± 0.65 | 1.19 ± 1.42 | 0.25 ± 0.67 |
| Pain | 0.23 ± 0.52 | 0.00 ± 0.00 | 0.65 ± 1.06 | 0.02 ± 0.14 | 0.79 ± 1.27 | 0.02 ± 0.14 | 1.00 ± 1.03 | 0.13 ± 0.44 |
| Redness | 0.27 ± 0.45 | 0.00 ± 0.00 | 0.50 ± 0.80 | 0.21 ± 0.82 | 0.54 ± 0.62 | 0.04 ± 0.29 | 0.44 ± 0.50 | 0.21 ± 0.85 |
| Headache | 0.23 ± 0.69 | 0.06 ± 0.32 | 0.44 ± 0.99 | 0.15 ± 0.51 | 0.52 ± 0.95 | 0.13 ± 0.64 | 0.40 ± 0.89 | 0.21 ± 0.68 |
| Fatigue | 0.60 ± 0.96 | 0.25 ± 0.64 | 1.15 ± 1.29 | 0.25 ± 0.79 | 1.21 ± 1.30 | 0.25 ± 0.79 | 0.94 ± 1.23 | 0.23 ± 0.72 |
| Concentration | 0.71 ± 0.99 | 0.17 ± 0.48 | 0.90 ± 1.31 | 0.13 ± 0.39 | 0.98 ± 1.16 | 0.13 ± 0.33 | 0.65 ± 1.18 | 0.15 ± 0.41 |
| Nervousness | 0.19 ± 0.57 | 0.00 ± 0.00 | 0.27 ± 0.68 | 0.00 ± 0.00 | 0.19 ± 0.39 | 0.00 ± 0.00 | 0.17 ± 0.56 | 0.02 ± 0.14 |
| Sleep Problems | 0.15 ± 0.46 | 0.08 ± 0.35 | 0.35 ± 1.08 | 0.08 ± 0.35 | 0.19 ± 0.57 | 0.08 ± 0.40 | 0.13 ± 0.61 | 0.02 ± 0.14 |

Side effects were reported on a scale ranging between 0 and 5 (larger numbers indicate more prominent side effects). Shown are group means and SD in young and elderly participants in different stimulation conditions.

**Table 2.3: ANOVA Results of the Side-Effects during tDCS with the within-subject factor tDCS Condition, and the between-subject factor Age Group.**

| *Variables* | *d.f.* | *d.f.$_{error}$* | *F-value* | $\eta_p^2$ | *p-value* |
|---|---|---|---|---|---|
| **Visual Phenomena** | | | | | |
| tDCS Condition | 3 | 282 | 3.588 | .037 | **.014*** |
| Age Group | 1 | 94 | 5.070 | .051 | **.027*** |
| tDCS Condition X Age Group | 3 | 282 | 1.145 | .012 | .331 |
| **Itching** | | | | | |
| tDCS Condition | 3 | 282 | 11.194 | .106 | **<.001*** |
| Age Group | 1 | 94 | 67.285 | .417 | **<.001*** |
| tDCS Condition X Age Group | 3 | 282 | 4.768 | .048 | **.003*** |
| **Tingling** | | | | | |
| tDCS Condition | 2.713 | 254.996 | 14.820 | .136 | **<.001*** |
| Age Group | 1 | 94 | 55.512 | .371 | **<.001*** |
| tDCS Condition X Age Group | 2.713 | 254.996 | 6.150 | .061 | **.001*** |
| **Burning** | | | | | |
| tDCS Condition | 3 | 282 | 4.329 | .044 | **.005*** |
| Age Group | 1 | 94 | 24.268 | .205 | **<.001*** |
| tDCS Condition X Age Group | 3 | 282 | 1.601 | .017 | .189 |
| **Pain** | | | | | |
| tDCS Condition | 2.616 | 245.879 | 7.066 | .070 | **<.001*** |
| Age Group | 1 | 94 | 50.477 | .349 | **<.001*** |
| tDCS Condition X Age Group | 2.616 | 245.879 | 4.086 | .042 | **.010*** |

Each item was analysed separately with a repeated-measures ANOVA. Mauchly's test of sphericity was conducted for each ANOVA, and Greenhouse–Geisser corrections were applied where sphericity assumptions were not met. Asterisks indicate significant effects ($p<.05$). d.f. = degrees of freedom; η2$p$ = partial eta squared.

**Table 2.4: ANOVA Results of the Side-Effects after tDCS with the within-subject factor tDCS Condition, and the between-subject factor Age Group.**

| *Variables* | *d.f.* | *d.f.<sub>error</sub>* | *F-value* | $\eta_p^2$ | *p-value* |
|---|---|---|---|---|---|
| **Redness** | | | | | |
| tDCS Condition | 2.407 | 226.257 | 3.538 | .036 | **.023*** |
| Age Group | 1 | 94 | 14.101 | .130 | **<.001*** |
| tDCS Condition X Age Group | 2.407 | 226.257 | 1.369 | .014 | .256 |
| **Headache** | | | | | |
| tDCS Condition | 2.743 | 257.822 | 1.621 | .017 | .189 |
| Age Group | 1 | 94 | 6.349 | .063 | **.013*** |
| tDCS Condition X Age Group | 2.743 | 257.822 | .690 | .007 | .547 |
| **Fatigue** | | | | | |
| tDCS Condition | 3 | 282 | 2.502 | .026 | .060 |
| Age Group | 1 | 94 | 27.998 | .229 | **<.001*** |
| tDCS Condition X Age Group | 3 | 282 | 2.468 | .026 | .062 |
| **Difficulty Concentrating** | | | | | |
| tDCS Condition | 3 | 282 | .818 | .009 | .485 |
| Age Group | 1 | 94 | 32.990 | .260 | **<.001*** |
| tDCS Condition X Age Group | 3 | 282 | 1.230 | .013 | .299 |
| **Nervousness** | | | | | |
| tDCS Condition | 2.635 | 247.726 | .496 | .005 | .661 |
| Age Group | 1 | 94 | 9.799 | .094 | **.002*** |
| tDCS Condition X Age Group | 2.635 | 247.726 | .786 | .008 | .488 |
| **Sleep Problems** | | | | | |
| tDCS Condition | 2.102 | 197.596 | 1.777 | .019 | .170 |
| Age Group | 1 | 94 | 2.749 | .028 | .101 |
| tDCS Condition X Age Group | 2.102 | 197.596 | 1.008 | .011 | .370 |

Each item was analysed separately with a repeated-measures ANOVA. Mauchly's test of sphericity was conducted for each ANOVA, and Greenhouse–Geisser corrections were applied where sphericity assumptions were not met. Asterisks indicate significant effects ($p<.05$). d.f. = degrees of freedom; η2$p$ = partial eta squared.

**Table 2.5: Exploring Age-Dependent Effects of Side Effects on Selective Sequence Learning via Linear Mixed Models**

| *Variables* | *d.f.* | *d.f.$_{error}$* | *F-value* | *p-value* |
|---|---|---|---|---|
| **Young Adults** | | | | |
| Side Effects | 1 | 146.46 | 0.0067 | .935 |
| Session Number | 1 | 144.65 | 6.5321 | **.012*** |
| Side Effects * Session Number | 1 | 146.92 | 0.0065 | .936 |
| **Older Adults** | | | | |
| Side Effects | 1 | 183.79 | 1.3582 | .245 |
| Session Number | 1 | 145.78 | 1.8689 | .174 |
| Side Effects * Session Number | 1 | 175.87 | 2.6635 | .105 |

The mixed model included the sum of side effect scores during tDCS (visual phenomena, itching, tingling, burning, and pain on a scale of 0-5 wherein larger numbers indicate more prominent side effects) and the session number (0-4). The dependent variable was the selective sequence learning magnitude (Baseline-standardised Reaction Time of Block 5 - Block 6). Asterisks indicate significant effects ($p<.05$). d.f. = degrees of freedom.
